## Supplementary figures and images for "Functional characterization of a “plant-like” HYL1 homolog in the cnidarian *Nematostella vectensis* indicates a conserved involvement in microRNA biogenesis"

### Figure_2_Source _data_1.pdf

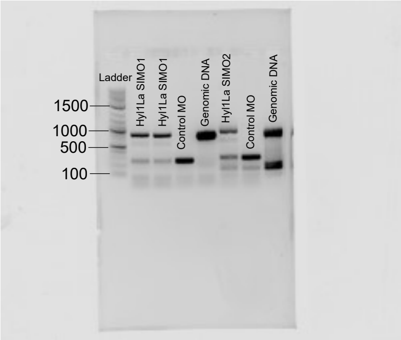

### Figure_2_Source _data_1.tif

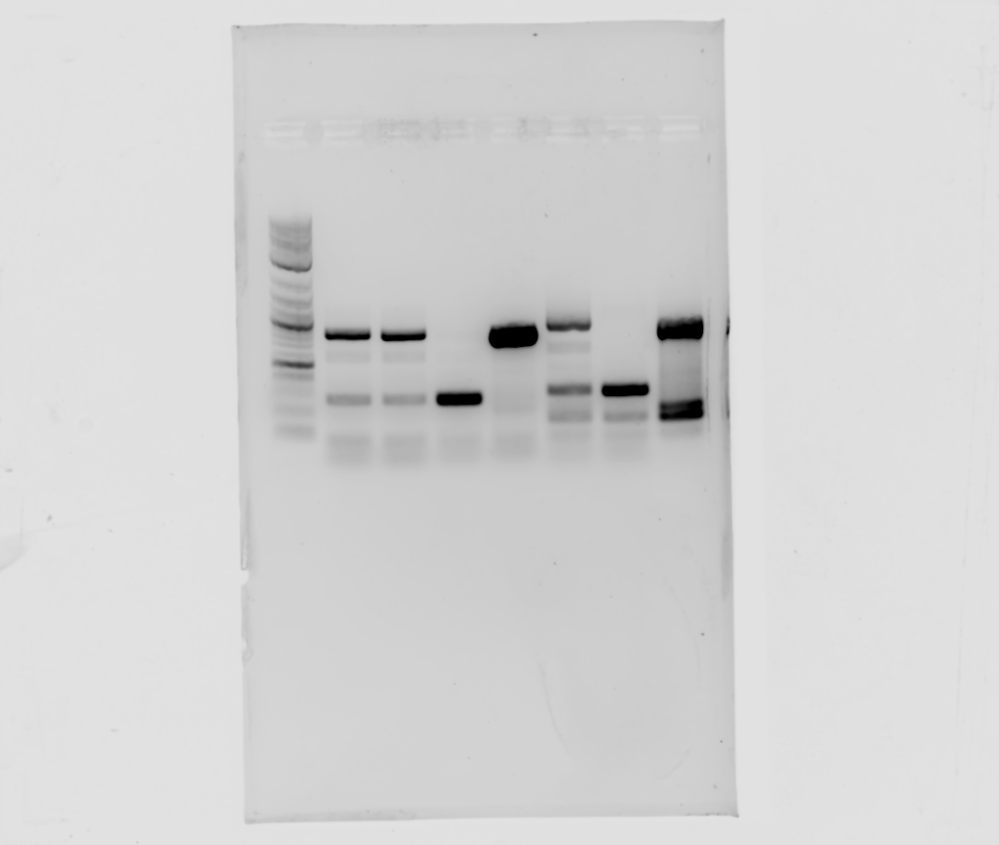

### Figure_5_Source _data_1.pdf

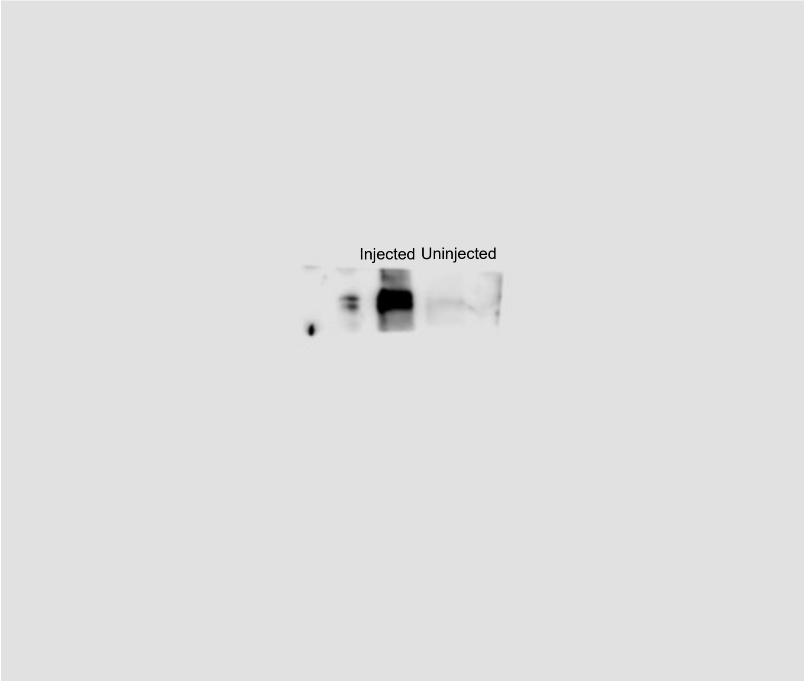

### Figure_5_Source _data_1.tif

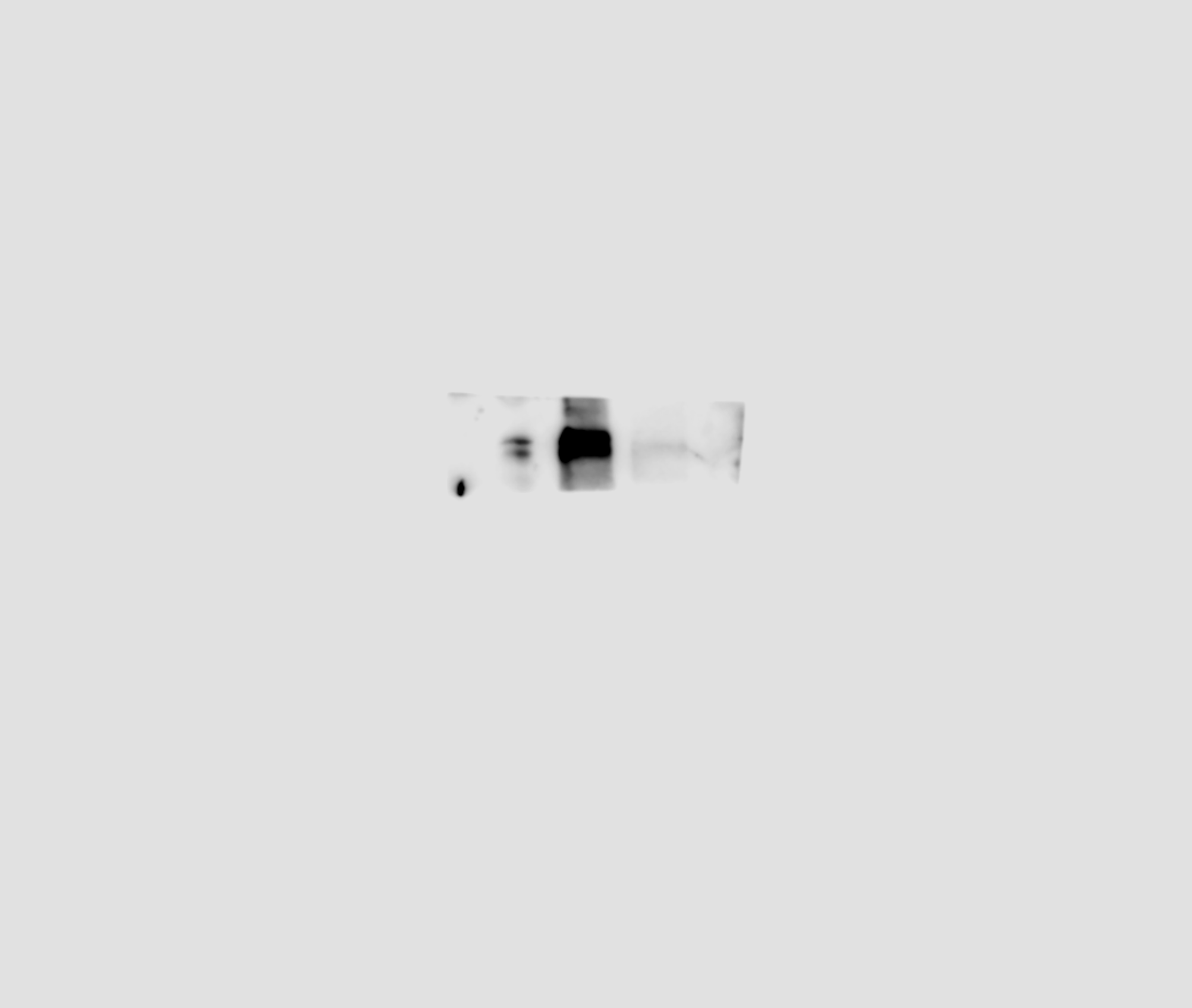

### Figure_5_Source _data_2.pdf

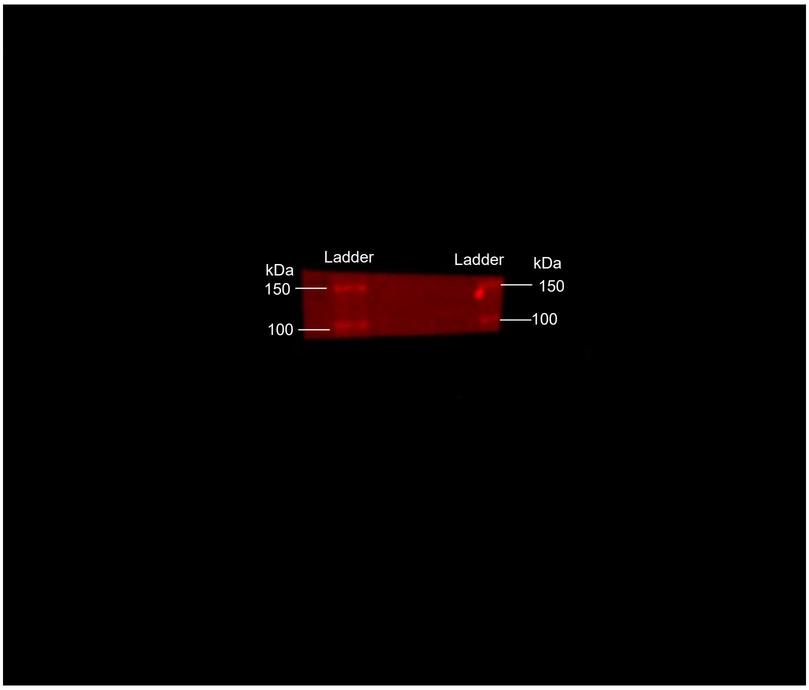

### Figure_5_Source _data_2.tif

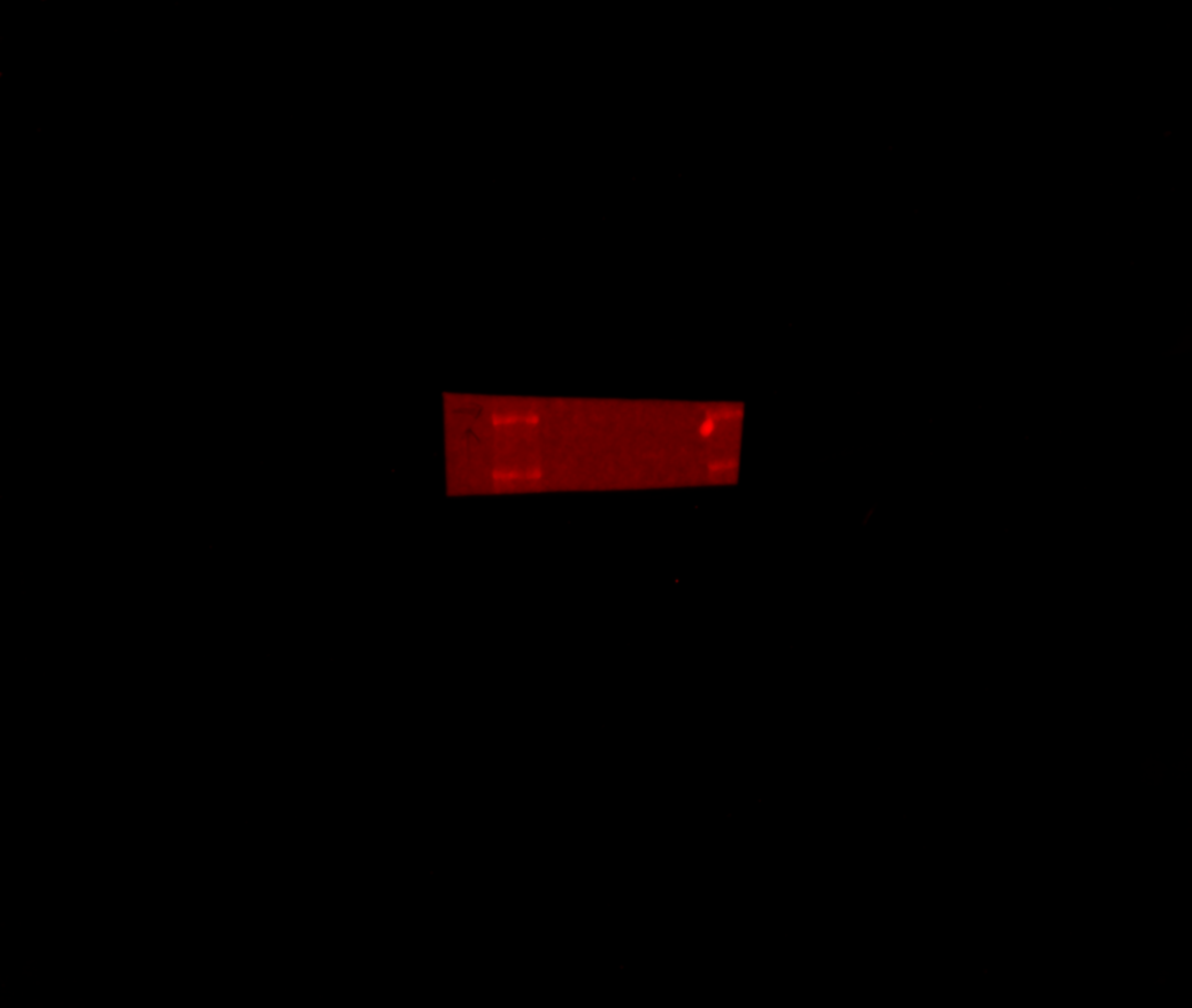

### Figure_5_Source _data_3.pdf

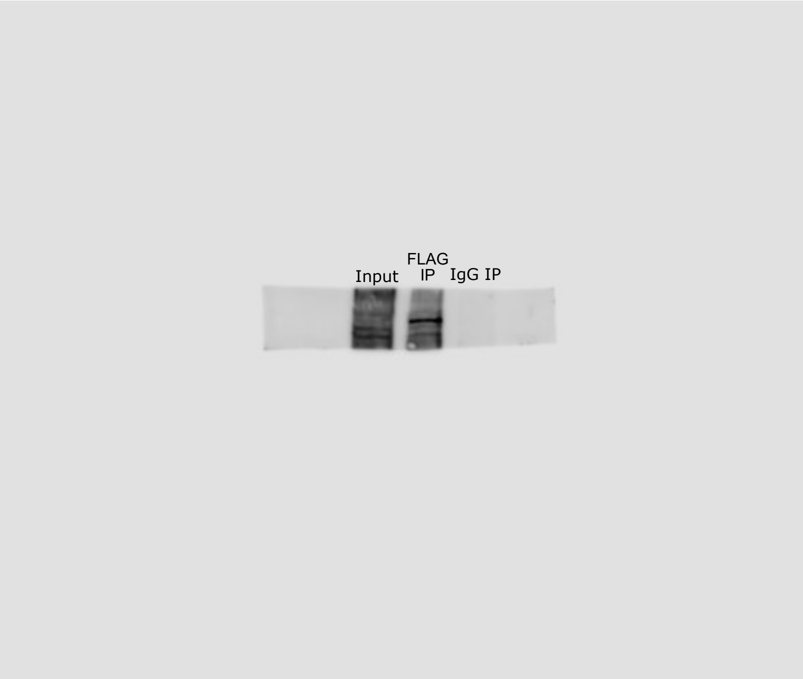

### Figure_5_Source _data_3.tif

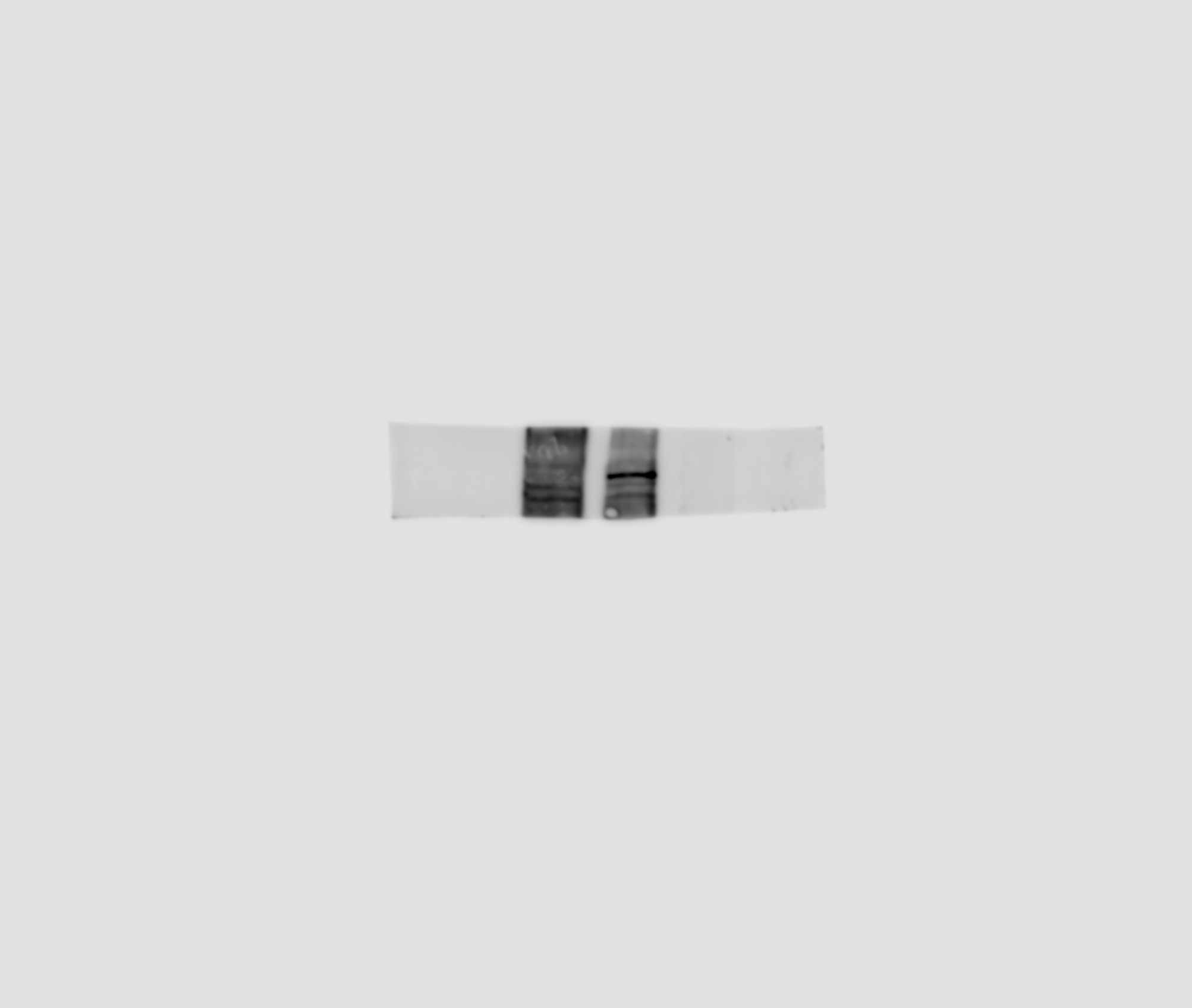

### Figure_5_Source _data_4.pdf

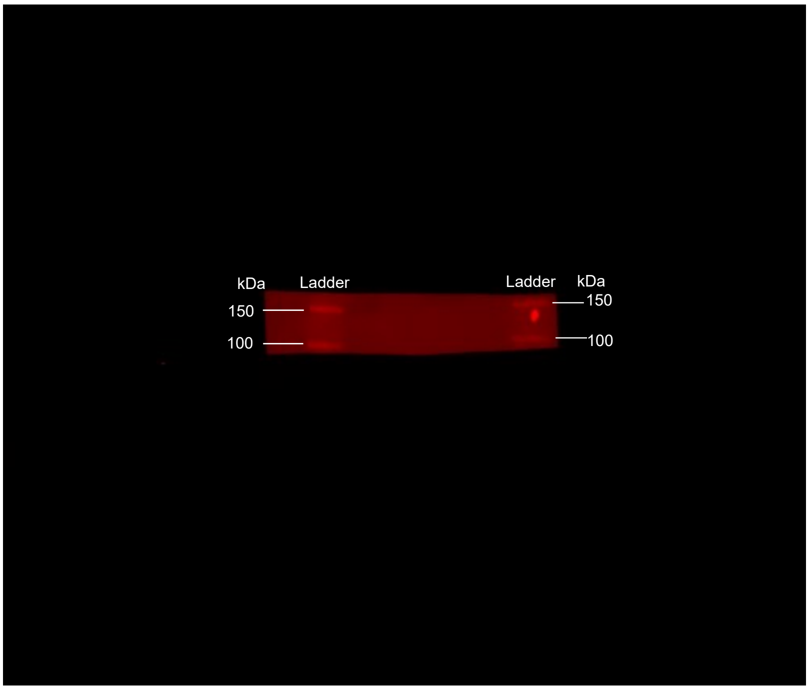

### Figure_5_Source _data_4.tif

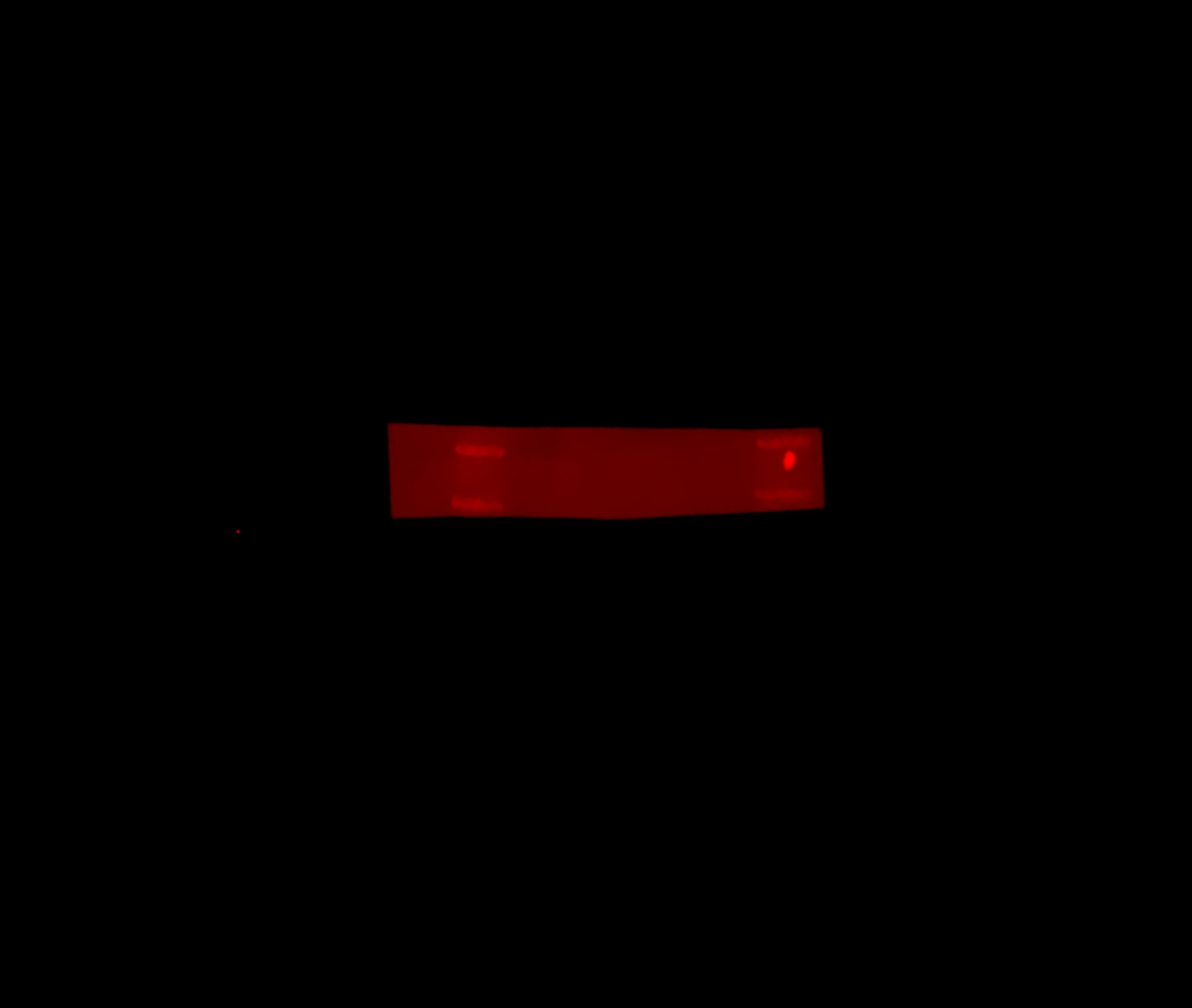

### Figure_5_Source _data_5.pdf

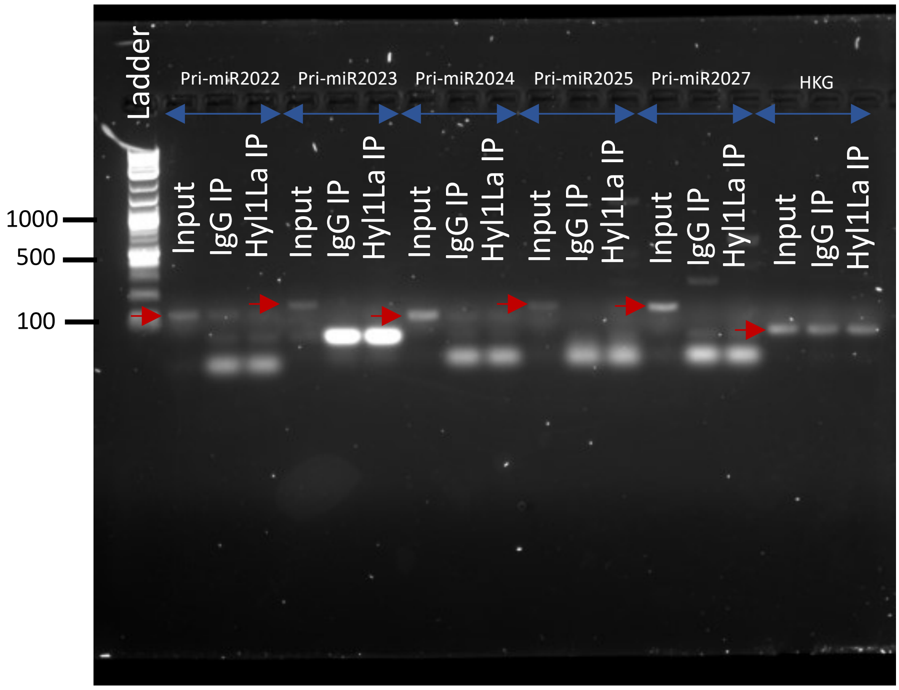

### Figure_5_Source _data_5.tif

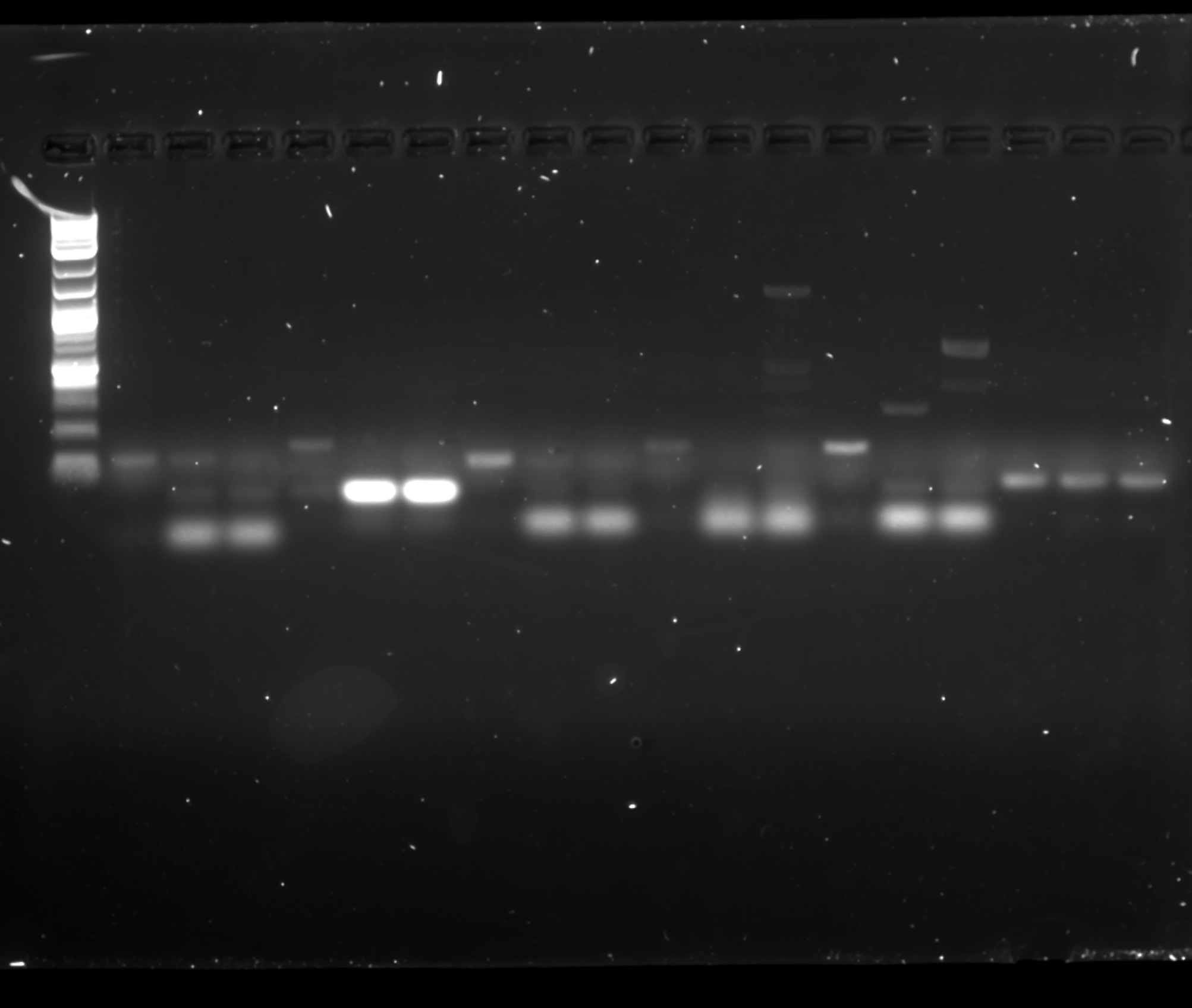

### Figure_5_Source _data_6.pdf

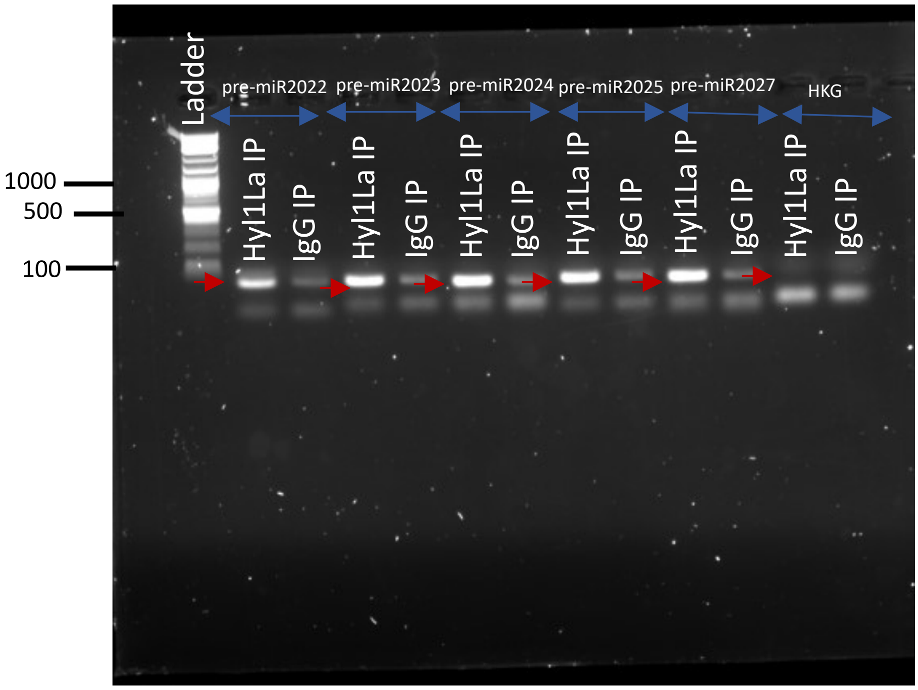

### Figure_5_Source _data_6.tif

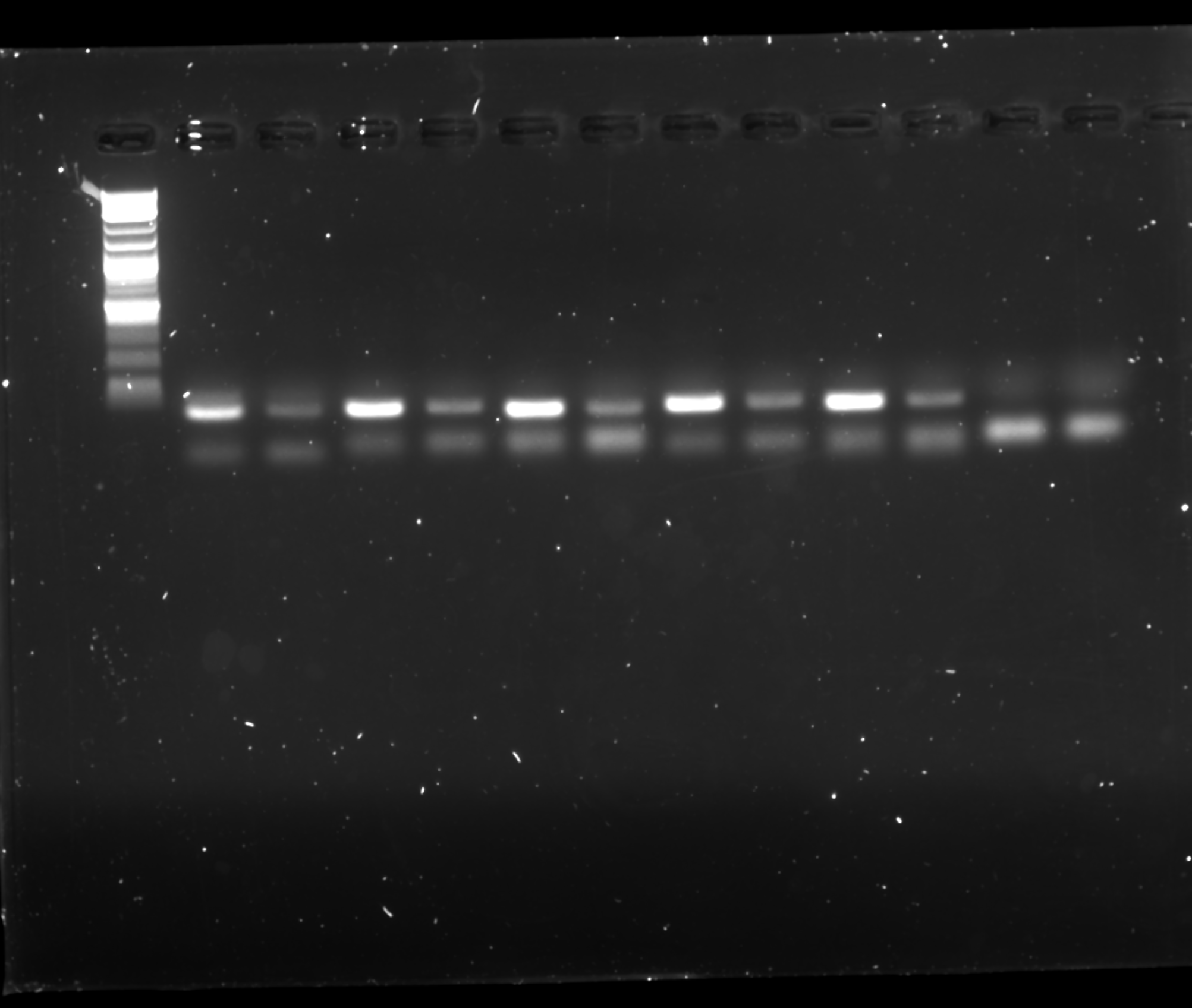

### Figure_6_Source _data_1.pdf

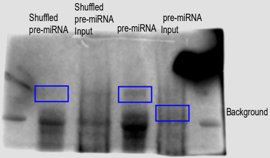

### Figure_6_Source _data_1.tif

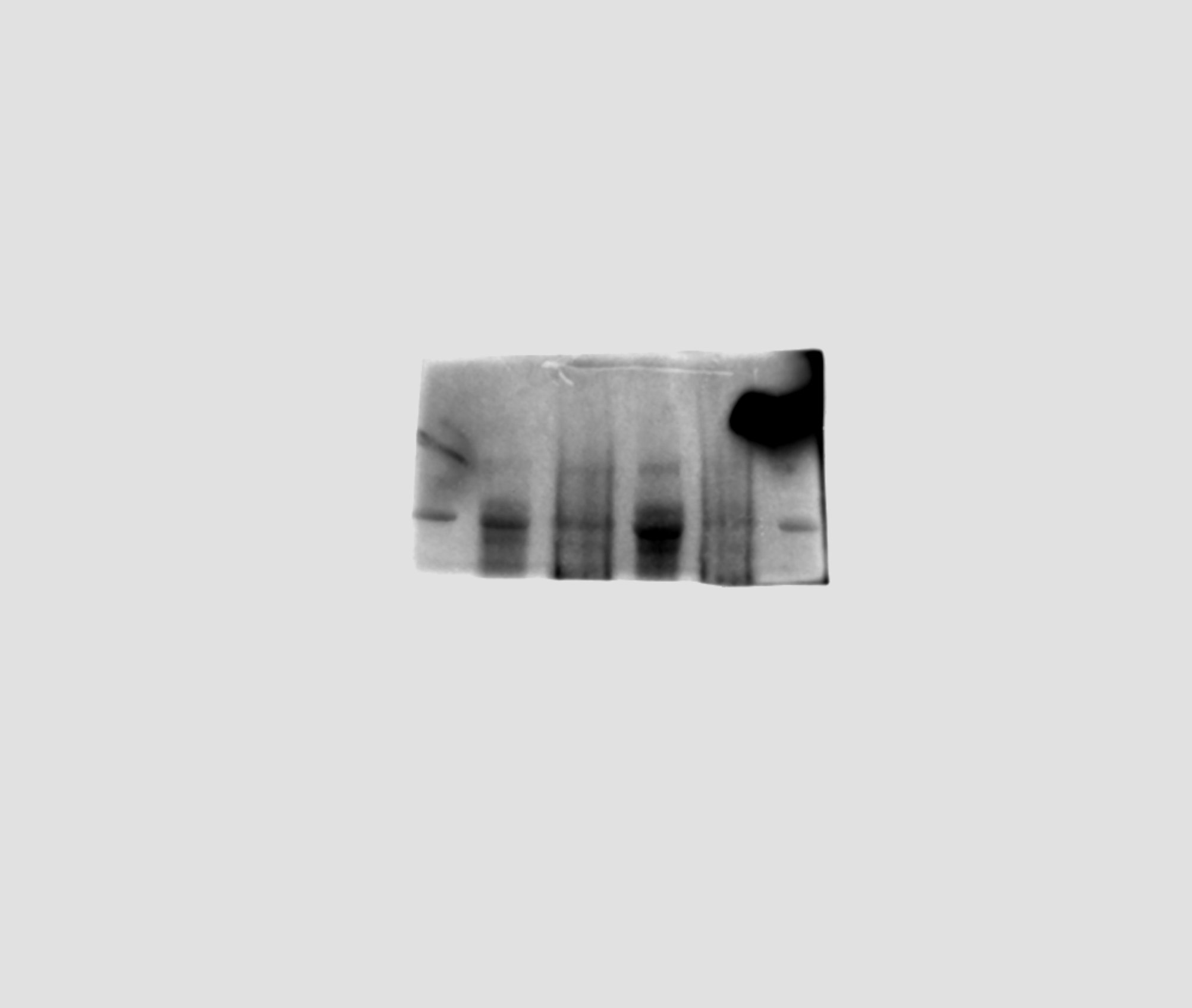

### Figure_6_Source _data_2.pdf

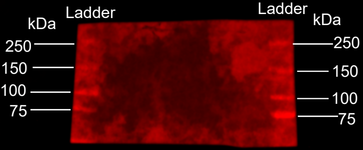

### Figure_6_Source _data_2.tif

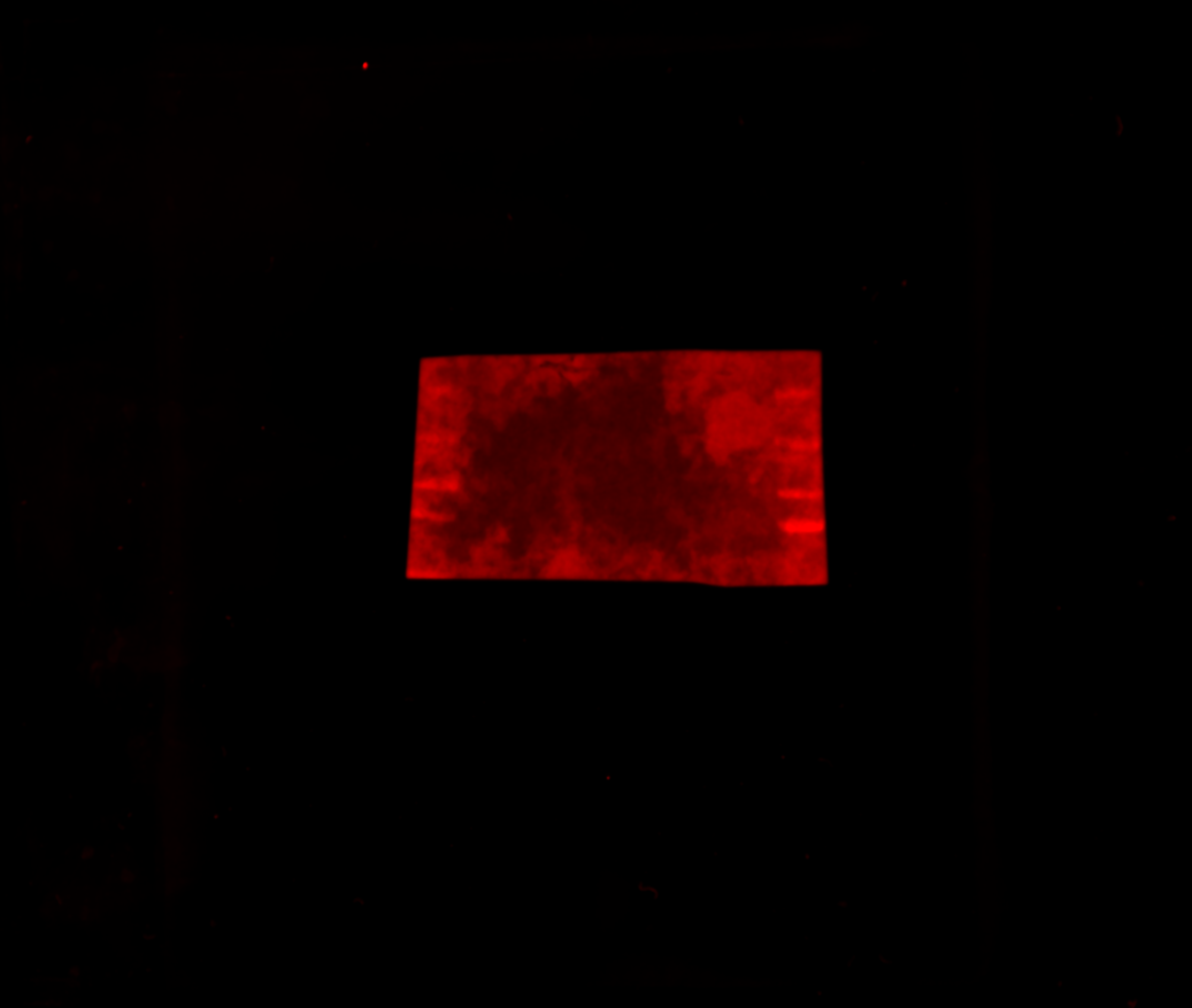

### Figure_6_Source _data_3.pdf

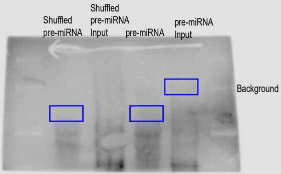

### Figure_6_Source _data_3.tif

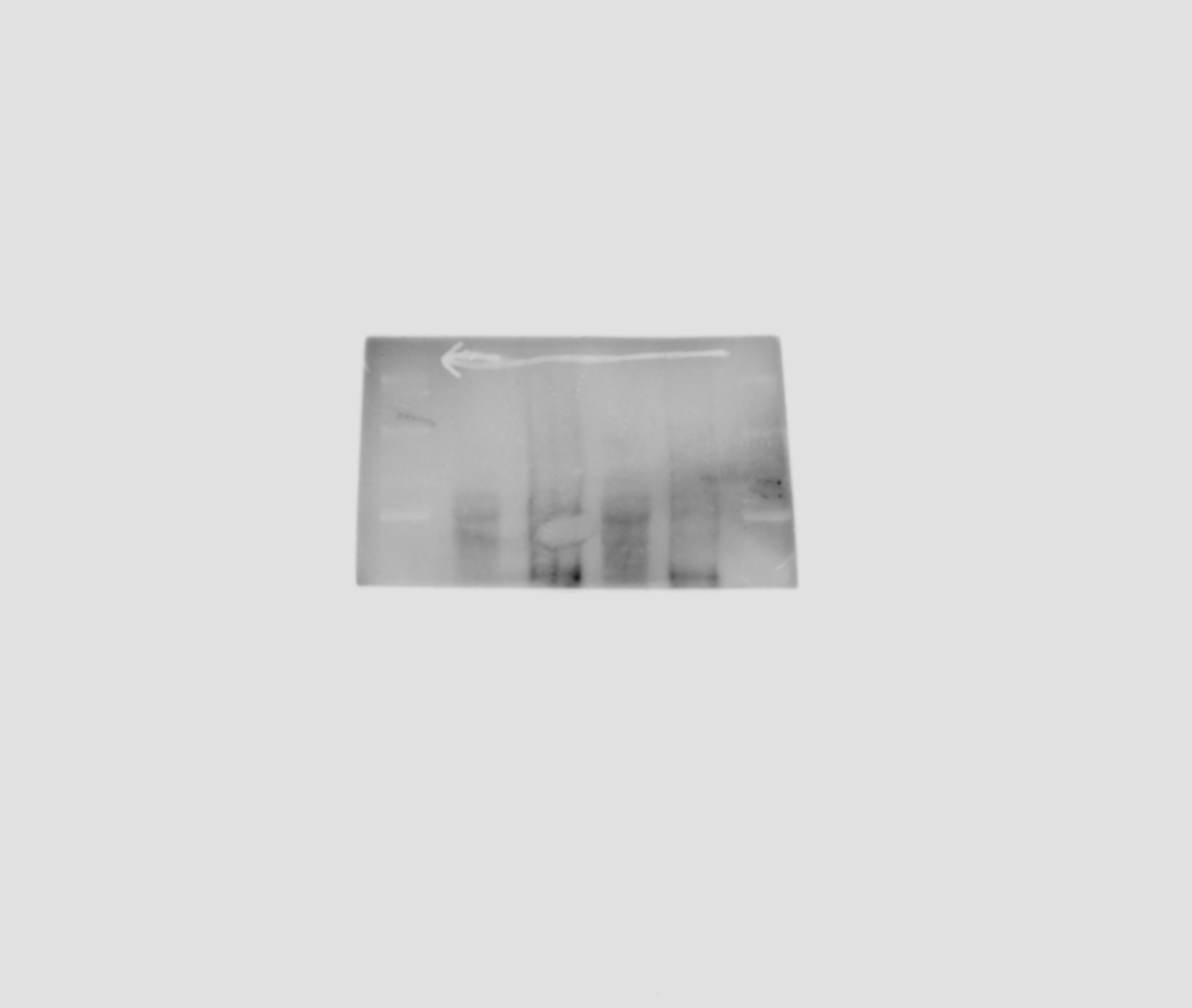

### Figure_6_Source _data_4.pdf

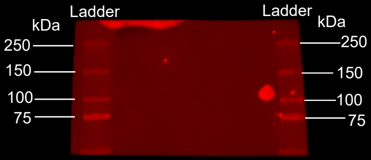

### Figure_6_Source _data_4.tif

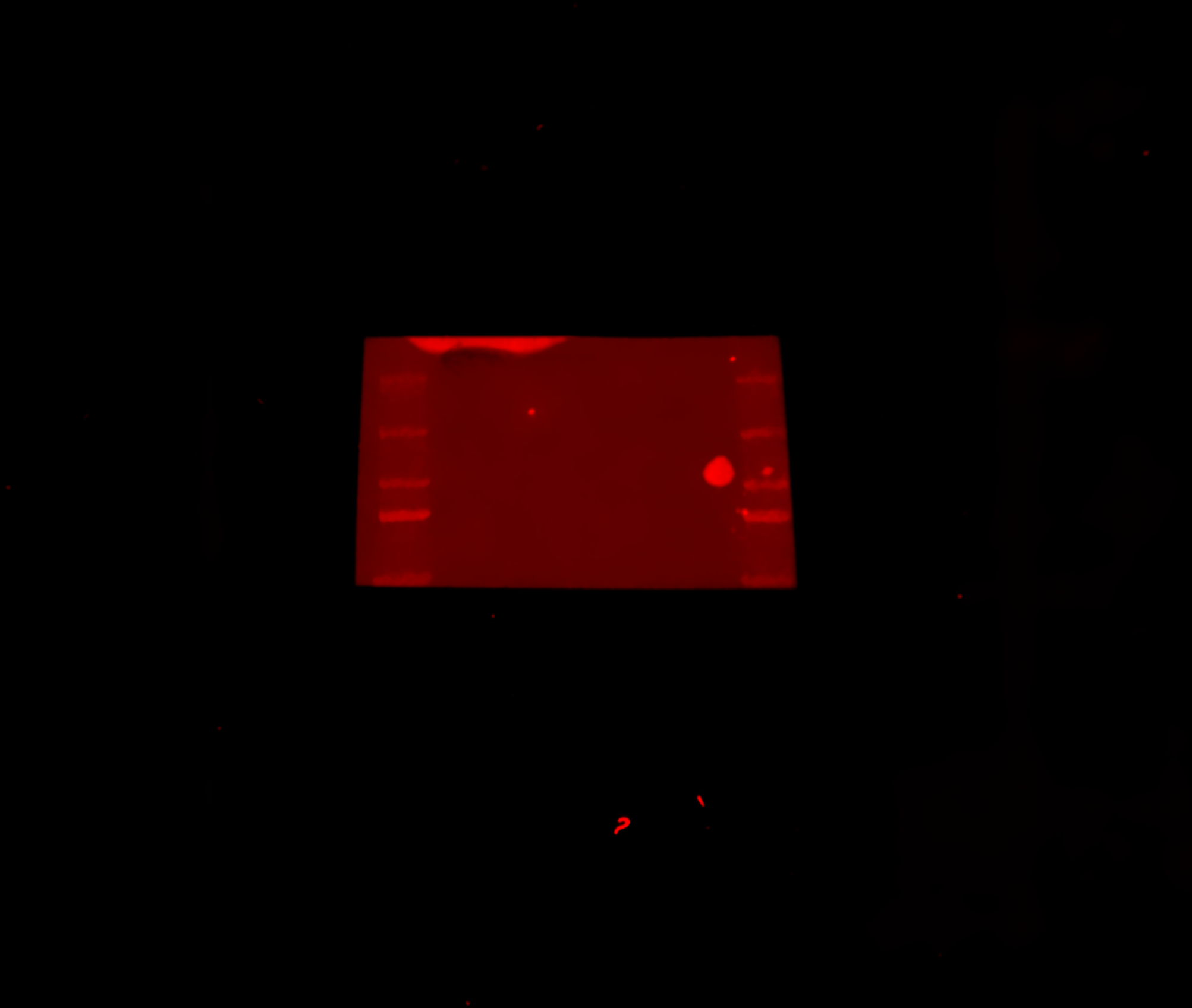

### Figure_6_Source _data_5.pdf

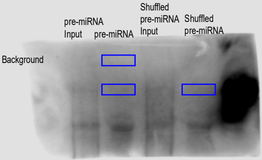

### Figure_6_Source _data_5.tif

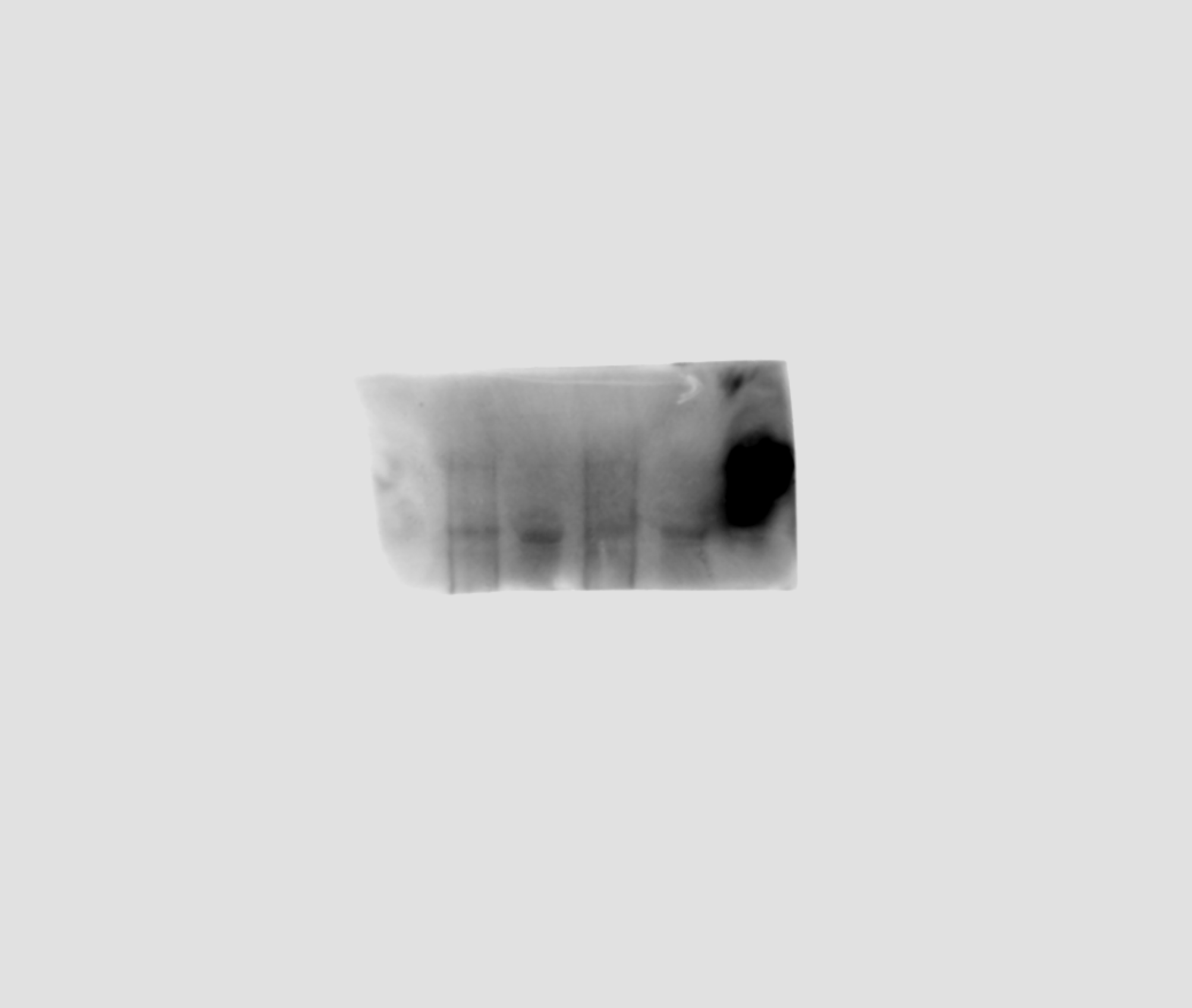

### Figure_6_Source _data_6.pdf

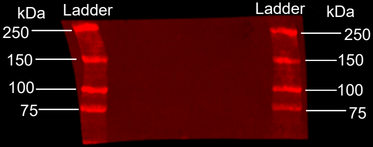

### Figure_6_Source _data_6.tif

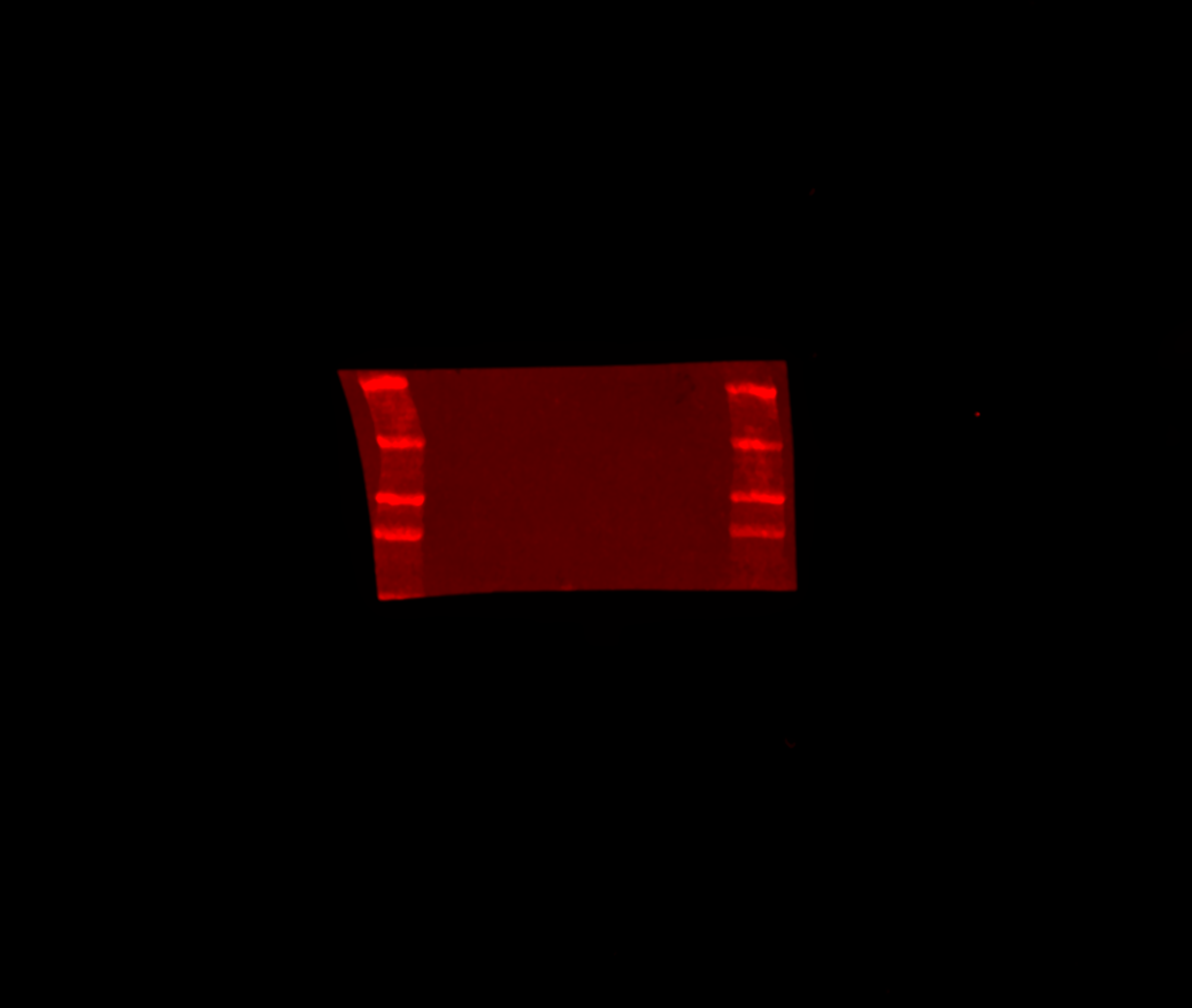
